# Top-down modulation and cortical-AMG/HPC interaction in familiar face processing

**DOI:** 10.1101/2022.04.14.488206

**Authors:** Xiaoxu Fan, Qiang Guo, Xinxin Zhang, lingxia Fei, Sheng He, Xuchu Weng

## Abstract

Humans can accurately recognize familiar faces in only a few hundred milliseconds, but the underlying neural mechanism remains unclear. Here we recorded intracranial electrophysiological signals from ventral temporal cortex (VTC), superior/middle temporal cortex (STC/MTC), medial parietal cortex (MPC) and amygdala/hippocampus (AMG/HPC) in 20 epilepsy patients while they viewed faces of famous people and strangers as well as common objects. In posterior VTC and MPC, familiarity-sensitive responses emerged significantly later than initial face-selective responses, suggesting that familiarity enhances face representations after they are first being extracted. Moreover, viewing famous faces increased the coupling between cortical areas and AMG/HPC in multiple frequency bands. These findings imply that the top-down modulation in local face-selective response and interactions between cortical face areas and AMG/HPC contribute to the superior recognition of familiar faces.

**Teaser:** Top-down modulation and cortical-AMG/HPC interactions contribute to the superior processing of familiar faces.

## Introduction

Recognizing familiar faces is remarkably accurate and fast in humans [1,2,3,4,5,6], even after not having seen them for a long time. Familiarity modulation has been extensively reported in face-processing system and regions involved in spontaneous activation of person knowledge, including medial temporal lobe structures (perirhinal cortex, amygdala, hippocampus) [7,8,9,10,11,12,13,14], anterior temporal regions [7,15,16,17,18,19,20,21,22], lateral aspect of temporal lobe [10,22], posterior cingulate gyrus [7,14, 23] and precuneus [22,23,24]. However, the neural mechanisms underlying the optimized processing of familiar faces remain largely unknown.

One hypothesis is that long-term familiarization tunes the bottom-up detectors to facial features and therefore makes familiar face processing more robust [1,25]. This theory predicts the very fast enhancement of face representation, at the stage of initial facial feature detection. Consistent with this hypothesis, identity and gender representations of familiar faces are enhanced very early, only a few milliseconds after they are extracted [26]. Eye tracking and psychophysics studies also identified ultrafast saccades to familiar faces and faster detection of familiar face even without focused attention and awareness, suggesting a feedforward familiarity processing mechanism [1,27]. However, it is still highly inconsistent across studies whether the N170 component of face-evoked ERP, which is often considered to represent the early stage of face processing, is modulated by familiarity [28,29,30,31,32,33,34,35,36].

Moreover, in most studies, only weak or even no fMRI activation differences between familiar and unfamiliar faces were reported in the core face areas [7,21,22,23,24,37] (occipital face area (OFA), fusiform face area (FFA) and posterior superior temporal sulcus (pSTS) [38]). Therefore, another theory proposed that face familiarity arises from activation of person knowledge and memory after detailed analysis of facial features, followed by top-down modulation of face representations after the initial response [39,40,41]. It has been postulated that the amygdala, hippocampus and adjacent cortical structures in the medial temporal lobe (MTL) interact with face network and have a top-down effect on areas responsible for face processing [75, 76].

However, the role and dynamics of interactions between the face network to both amygdala and hippocampus are still unclear.

In our view, the key to resolve the conflicting hypotheses is to have concrete and precise data on whether familiarity enhancement of face representation in core-face selective regions occur at an initial or late stages of face processing. In the human brain, arguably the intracranial electroencephalography data provide the best resolution both spatially and temporally. We thus recorded stereo-electroencephalography (sEEG) signals from the ventral temporal cortex (VTC), superior/middle temporal cortex (STC/MTC), medial parietal cortex (MPC) and amygdala/hippocampus (AMG/HPC) in 20 pre-surgical epilepsy patients while they viewed images of famous and unfamiliar faces as well as objects. We observed familiarity sensitive responses in bilateral posterior VTC (pVTC), left posterior middle temporal cortex and left MPC. In pVTC and MPC, familiarity enhancement begins at about 240 ms and 450 ms after the presentation of faces, in other words, about 100 ms and 190 ms after the initial face-selective responses, respectively. Moreover, the functional connectivity between cortical areas and AMG/HPC was increased when participants viewed famous faces compared with unfamiliar faces. Our findings provide direct neurophysiological evidence in humans for the elevated involvement of top-down mechanisms in processing famous faces and further our understanding of the neural basis of familiar face recognition.

## Results

### Data collection and design

We performed direct intracranial recordings with depth electrodes in 20 pre-surgical epilepsy patients (14 males and 6 females, age 14-53 years old). Each electrode consists of 10– 18 finely spaced recording contacts. More electrodes were implanted in left than right hemisphere due to the clinical purposes of seizure localization. Participants were asked to do an oddball detection task while viewing images of famous faces, unfamiliar faces and objects (**Figure 1A**). We focused on signals recorded from VTC (407 contacts), STC/MTC (517 contacts), amygdala (51 contacts), hippocampus (39 contacts) and MPC (105 contacts) (**Figure 1B**).

**Figure 1.**
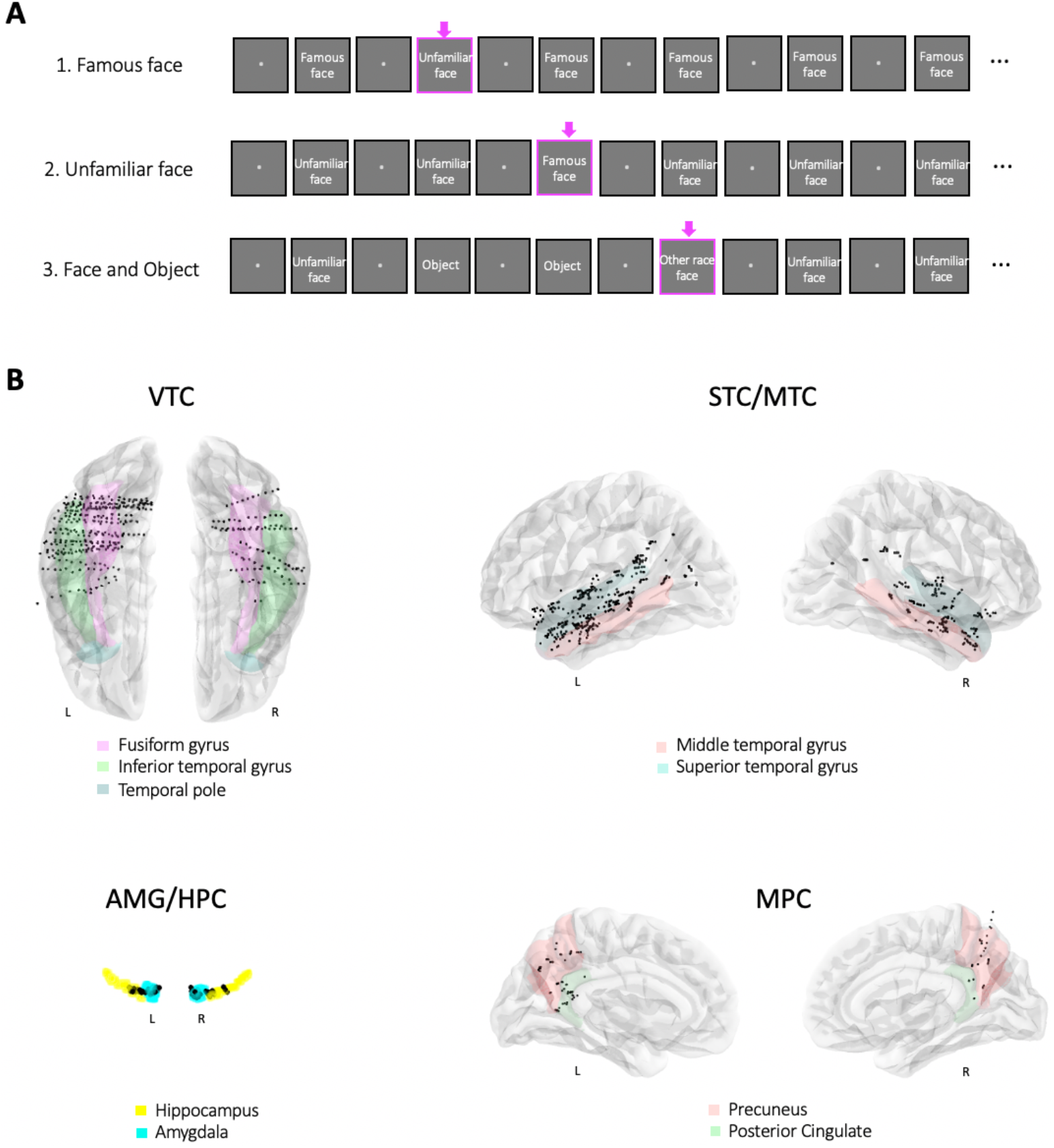
Experiment Design and Contact Locations. (**A**) The oddball paradigm. In session 1, rare unfamiliar faces were embedded in a sequence of frequent famous faces. In session 2, rare famous faces were embedded in a sequence of frequent unfamiliar faces. In session 3, rare other race faces were embedded in a sequence of frequent unfamiliar faces and objects. The oddball targets are marked by pink arrow. Pictures used in real experiment are shown as words to avoid the inclusion of photographs of people in the figure. (**B**) Contact locations across 20 participants displayed on a three-dimensional transparent brain (fsaverage in FreeSurfer) in the MNI space. Each black dot represents a single contact. The anatomical subdivisions are shown in corresponding colors. VTC, ventral temporal cortex; STC/MTC, superior/middle temporal cortex;

### Spatial Distribution of Familiarity-sensitive Responses

High-frequency broadband (HFB) signal from sEEG is a correlate of spiking activity of neurons adjacent to the recording site and thus currently is interpreted as a reflection of local neural activity [42,43,44,45,46,47]. In order to investigate the local engagement of the various brain areas, we analyzed HFB (110-140 Hz) responses at all contacts to each category of images (famous faces, unfamiliar faces and objects). Of all the recorded contacts, 30.21% (n=338) had significant HFB responses to at least one category of images relative to baseline and only these task-active contacts were included in the data analysis. Then a contact was considered as face-selective if it displayed significantly greater HFB power to faces than to objects between 100 and 700 ms after stimulus onset. Cluster-based permutation testing was applied to assess the significance of the response (faces > objects, 10000 permutations, p < 0.05 FDR corrected for the number of contacts within each brain area). Similarly, familiarity-sensitive contacts were identified as sites where significantly higher HFB responses were seen to the famous faces compared to unfamiliar faces.

Out of 338 task-active contacts, 13 contacts in pVTC (y <-45), 10 contacts in anterior VTC (aVTC, y>= -45), 7 contacts in posterior STC/MTC (pSTC/MTC, y <-45) and 5 contacts in MPC were face-selective (**Figure 2**). Eleven contacts in bilateral pVTC, 2 contacts in left pSTC/MTC and 9 contacts in left MPC showed stronger signals to famous faces compared to unfamiliar faces. The locations of 10 out of 13 contacts in the temporal cortex showing familiarity enhanced responses correspond to FFA and pSTS in the core face-processing network (**Table 1**)[38,48,49].

**Figure 2.**
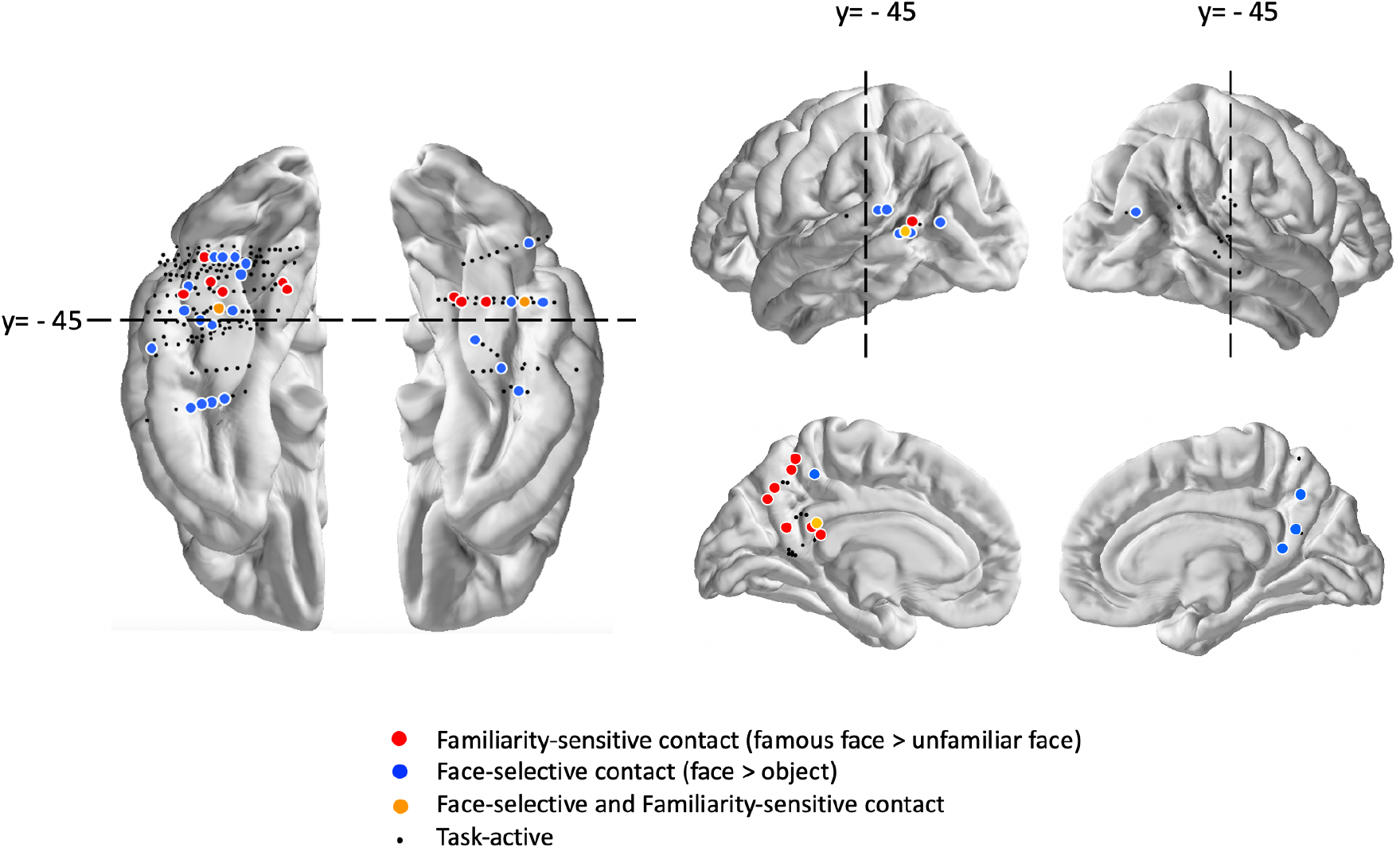
Spatial distribution of face-selective and familiarity-sensitive contacts. All task-active contacts from 20 participants are projected to cortical surface to facilitate visualization. Contacts that showed selectivity are displayed larger than their actual sizes.

**Table 1.**
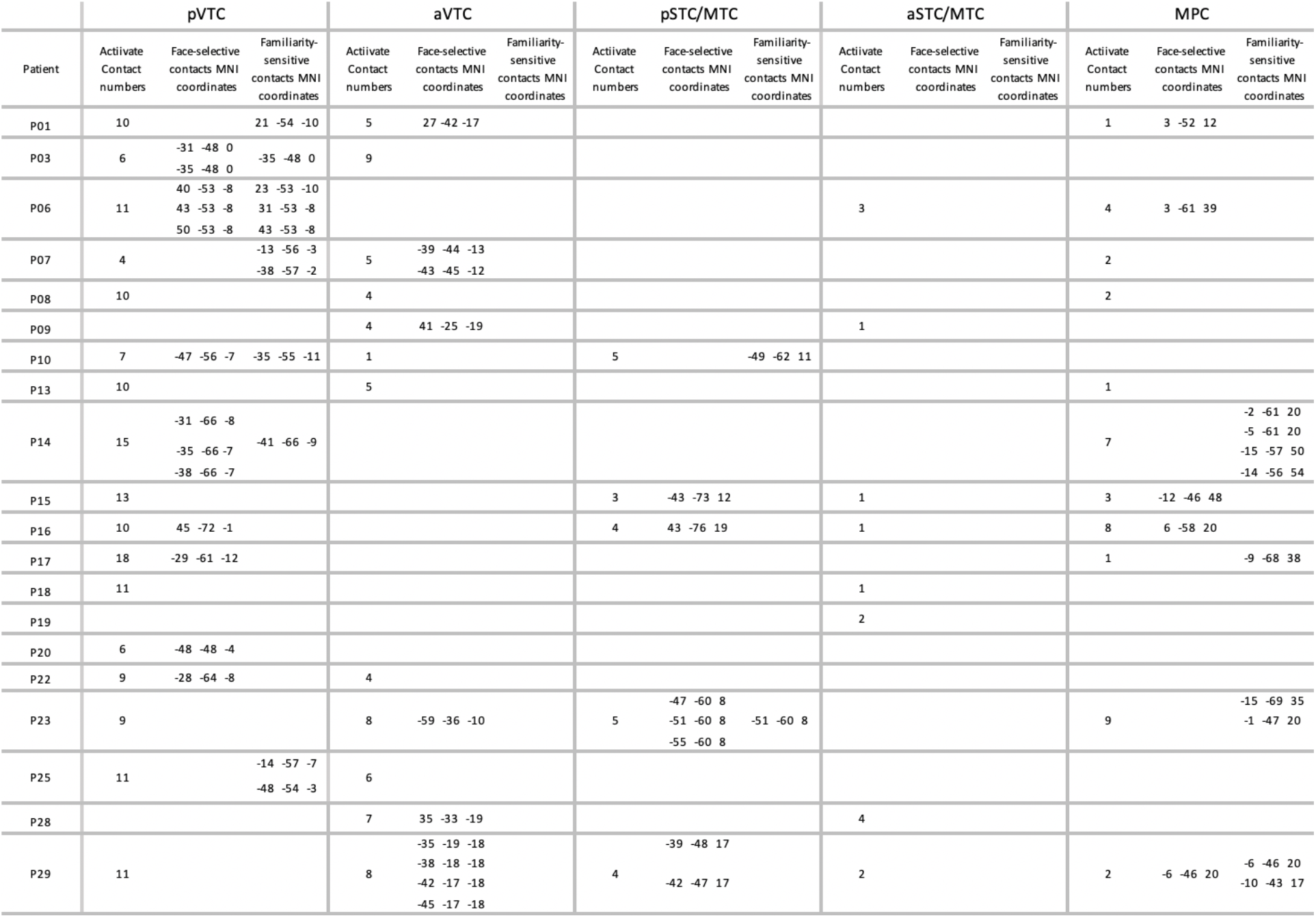
MNI coordinates of face-selective and familiarity sensitive contacts.

### Temporal dynamics of familiarity modulation

Following the identification of the face-selective and familiarity-sensitive contacts, we then investigated the temporal dynamics of HFB responses in familiarity-sensitive and face-selective contacts. Two contacts in fusiform gyrus, one contact in posterior middle temporal gyrus and one contact in posterior cingulate showed both face-selective and familiarity-sensitive responses. In all these contacts, the familiarity sensitivity emerged later than face selectivity (**Figure 3**).

**Figure 3.**
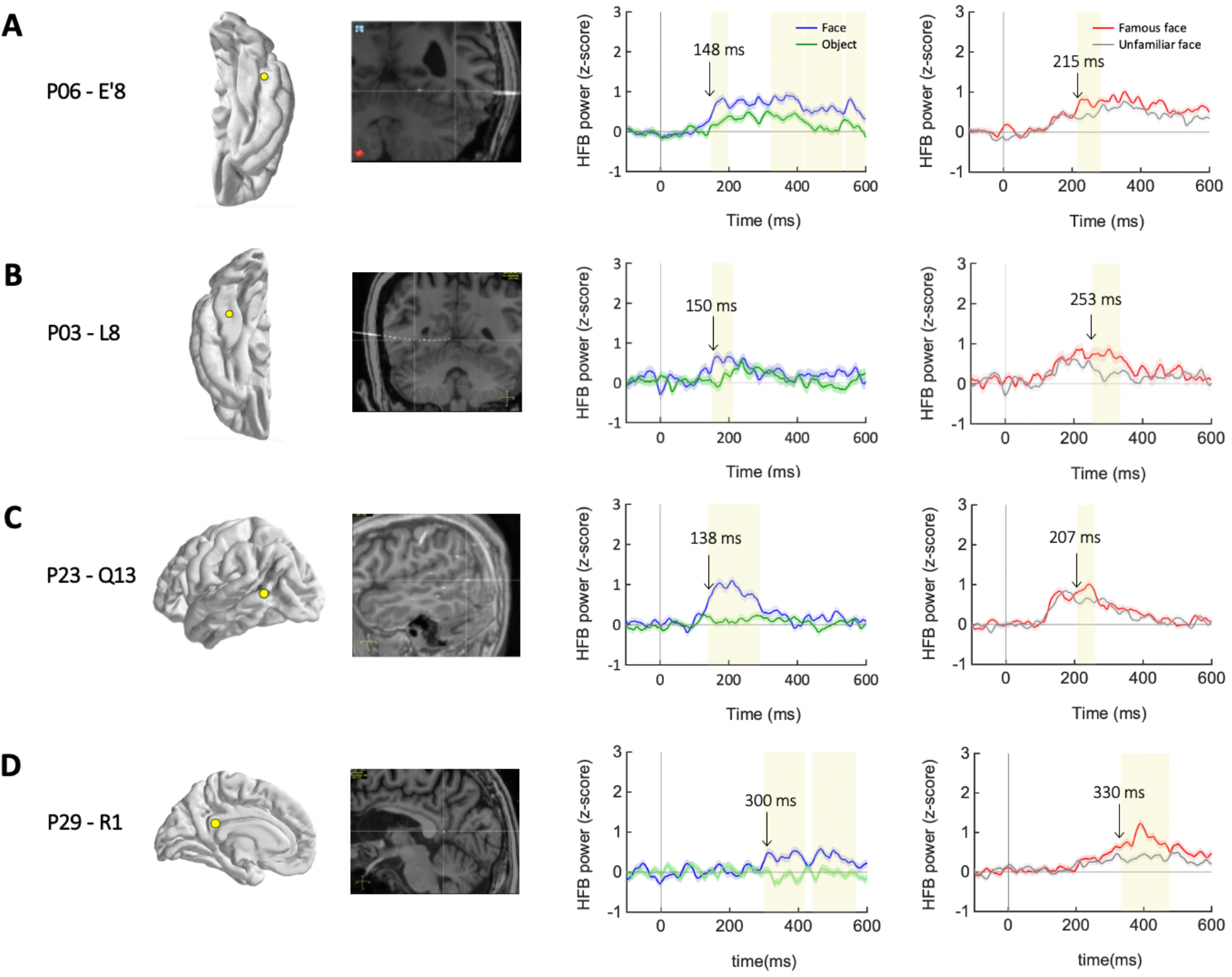
Locations and time courses of four representative contacts. Example contacts that showed both face-selective and familiarity-sensitive responses in 4 participants (P03, P06,P23 and P29) are displayed on cortical surface (yellow dot) and postoperative computed tomography (CT) images coregistered to preoperative MRI. z-scored HFB power at each contact in response to each category is colored blue (face), green (object), red (famous) and gray (unfamiliar). Yellow shaded area indicates the time clusters of significantly larger responses to famous than unfamiliar faces (Right) or face than object (Left) (permutation tests, p < 0.05). The onset latencies of face-selective or familiarity-sensitive responses are labeled above the time courses.

To further investigate the temporal relationship between face-selective and familiarity-sensitive responses, we averaged the individual time courses across all face-selective and familiarity-sensitive contacts respectively and the results are shown in **Figure 4**. Averaged time courses of famous vs. unfamiliar faces are not shown in pSTC/MTC and aVTC due to the limited number of familiarity-sensitive contacts. We compared the onset latency of face-selective and familiarity-sensitive responses at the population level. In pVTC, contacts begin to show face-selective responses as early as 125 ms after stimulus presentation (median, 125 ; interquartile range (IQR), 117-149), while familiarity-sensitive responses appeared 240 ms after stimulus presentation (median, 240 ; IQR, 188-463) (**Figure 4A** bottom). Familiarity-sensitive responses emerged significantly later than the onset of face-selective responses (Moods median test, p=0.004). In MPC, the onset latency of familiarity-sensitive response is 450 ms (median,450; IQR,380-473), which is also significantly later than face-selective responses (median, 260; IQR: 189-289)(Moods median test, p=0.005).

**Figure 4.**
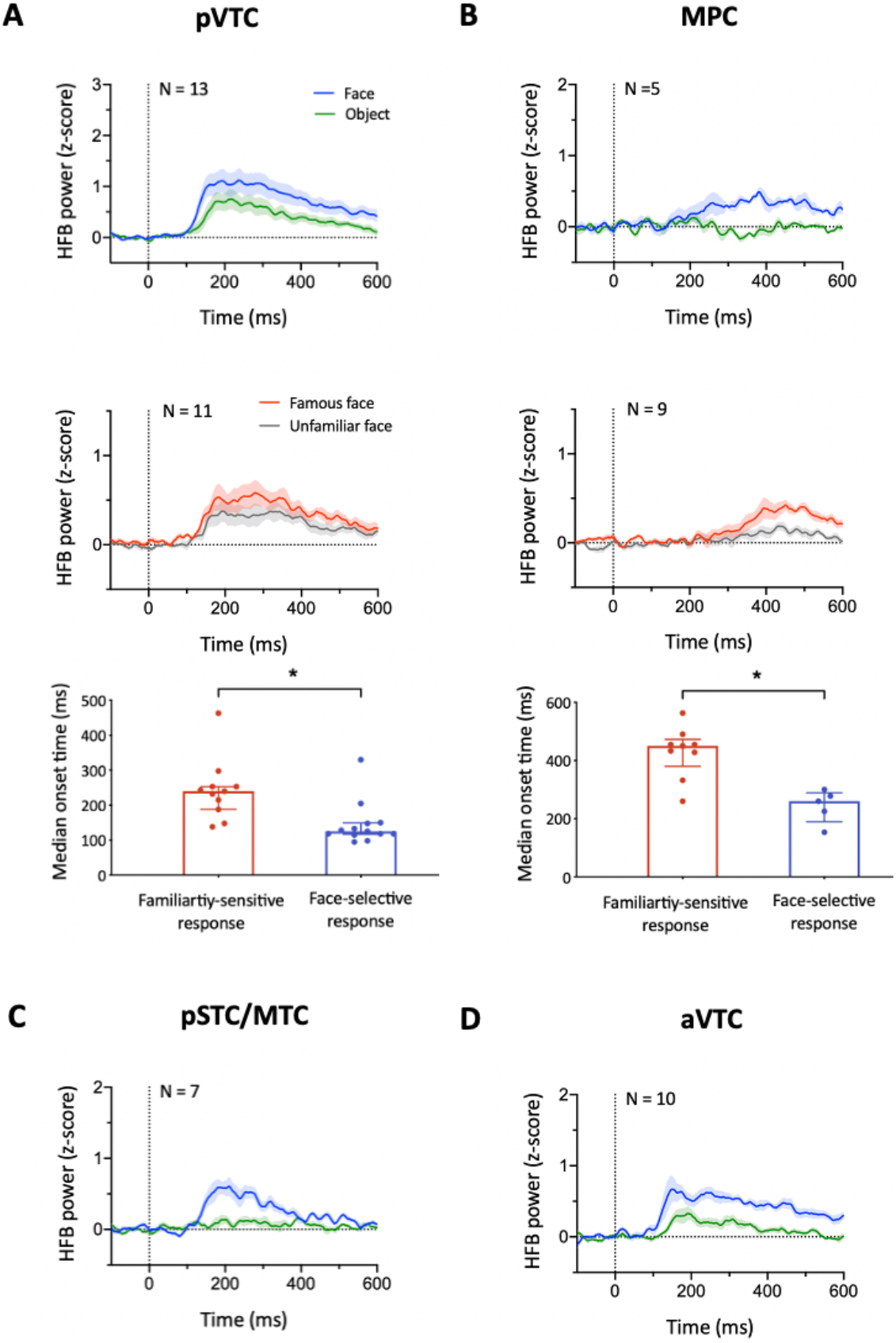
Population analysis of face-selective and familiarity-sensitive responses. (**A** and **B**) Average time course (mean ± SEM) of z-scored HFB power for each category in face-selective contact sites (Top) and familiarity-sensitive contact sites (Middle). N is the number of contacts. Onset latencies of face-selective and familiarity-selective responses are at the bottom (median + interquartile range(error bars)). *, p<0.05, Moods median test. (**C** and **D**) Average time course (mean ± SEM) of z-scored HFB power for each category in face-selective contact sites.

Our findings indicate that familiarity modifies face representation in bilateral pVTC and left MPC but such familiarity effect occurred significantly later than the initial process of face detection, suggesting the involvement of top-down modulation during familiar face processing.

### Familiarity modulation in Cortical - MTL connectivity

Structural and functional connectivity between cortical face processing areas and AMG/HPC have been identified in both human and monkey brains, and such connections were suggested to contribute to face-related memory functions [62,63,64,65]. Since famous faces in essence are well-remembered faces, presumably their processing would strongly engage the interactions between cortical face processing areas and AMG/HPC. To test this prediction, we evaluated the phase-locking value (PLV)[50,51] between cortical areas (pVTC, aVTC, pSTC/MTC, aSTC/MTC and MPC) and AMG/HPC in a subset of patients (n=17). Only task-active contact pairs within-patient were included in the calculation of PLV (pVTC-AMG/HPC: 159 pairs, aVTC-AMG/HPC: 68 pairs, pSTC/MTC-AMG/HPC: 26 pairs, aSTC/MTC-AMG/HPC:25 pairs, MPC-AMG/HPC: 66 pairs.). **Figure 5A** shows the temporal distribution of significantly greater PLV during famous face trials than unfamiliar face trials (cluster-based permutation test, Bonferroni corrected p < 0.05) in theta (3 - 5 Hz), alpha (8 - 12 Hz), beta (12 - 30 Hz)and gamma band (30 - 60 Hz). Viewing famous faces increased PLV between pVTC and AMG/HPC at both early stages (125-177 ms in α band; 109-159 ms in β band, 101-134 ms in γ band) and late stages (θ: 443-570 ms, α: 378-520 ms; β: 368-468 ms, γ: 393-457 ms) (**Figure 5B**). The increase of PLV for famous faces was also observed fairly late in aVTC-AMG/HPC pairs (β: 486-556 ms, γ: 473-546 ms), pSTC/MTC-AMG/HPC pairs (α: 535-574 ms) and MPC-AMG/HPC pairs (β: 501-563 ms, γ: 492-558 ms). These findings showed enhanced functional connectivity between cortical face processing areas and AMG/HPC during famous face processing. The early interaction between pVTC and AMG/HPC may support the prioritization of familiar face processing while the late interaction between AMG/HPC and more broad cortical areas likely support the additional processing of stored information related to famous faces.

**Figure 5.**
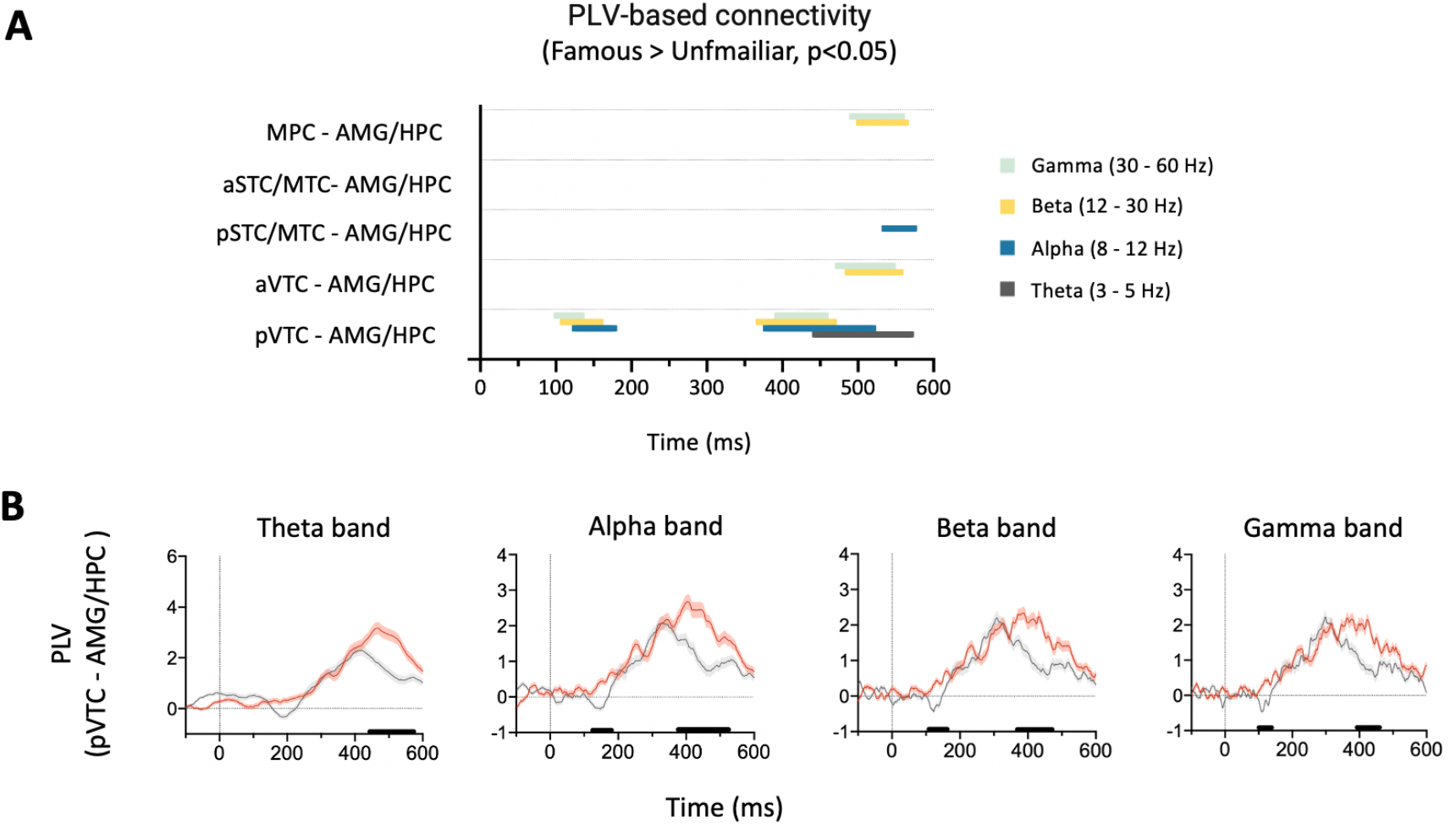
Phase locking value between cortical areas and AMG/HPC. (**A**) The durations of significant greater PLV induced by famous faces compared to unfamiliar faces between cortical-AMG/HPC pairs in theta, alpha, beta and gamma band. (**B**) Average time courses (mean+ SEM) of PLV changes from baseline across all pVTC-MTL pairs in theta, alpha, beta and gamma band. Black horizontal bars are the durations of significant difference between PLV of famous faces and unfamiliar faces.

Together, this study for the first time uncovered the temporal dynamics of familiar face processing in multiple face and memory-relevant areas in the human brain. Our results showed that famous face processing was associated with late enhancement of face representation in posterior temporal face areas and MPC as well as multiple stages of increased connectivity between these areas and AMG/HPC, indicating the involvement of top-down neural mechanisms for the efficient recognition of familiar faces and the and association of these faces with their stored information.

## Discussion

Using direct electrophysiological recording, our results provide important insights on the mechanisms underlying efficient familiar face recognition. We observed stronger signals in pVTC and MPC following the initial face response, and at multiple stages, enhanced interactions between cortical areas and AMG/HPC during famous faces processing. These findings support the hypothesis that repeated exposure with specific faces in daily life enable efficient retrieval of related person knowledge and in turn facilitate face processing in a top-down manner.

Functional neuroimaging studies have identified a series of brain regions that respond more strongly to familiar faces than to unfamiliar faces [7,22,38,52]. However, familiarity-dependent modulation in core face-processing regions was only reported in a few studies [7,21,22,23,24,37]. Further, fMRI studies were unable to provide precise timing information to elucidate whether the observed familiarity effect results from better tuning to the features in the initial processing or from top-down modulation following the initial general face processing. In the current study, the locations of 10 of the 13 familiarity-sensitive contacts identified in temporal lobe are consistent with FFA and pSTS. In pSTC/MTC, all familiarity-sensitive contacts were found in the left hemisphere, most likely because there were more electrodes placed in the left hemispheres among the patients we tested. Importantly, both individual results and group results clearly showed that the familiarity-enhancement effect followed the initial face category responses. Notably, the face-selective contacts and familiarity-sensitive contacts only partially overlapped in pVTC, pSTC/MTC and MPC. One possibility is that familiarity-related feedback to temporal cortex modulates the units engaged in feedforward processing of faces as well as new units in the nearby region [53]. Another possibility is that the HFB responses in many face-selective contacts were at or near their ceiling level responses and therefore could not be significantly increased by familiarity.

Effects of face familiarity have been observed in multiple areas outside the face-processing regions, for example, amygdala [7,11,14], anterior inferior temporal (AIT) cortex [7,10], orbitofrontal gyrus [7], and medial parietal cortex (MPC) [14,22,66]. In the human MTL (hippocampus, amygdala, entorhinal cortex and parahippocampal gyrus), subsets of neurons respond selectively to different pictures of given famous people [54]. These neurons with invariant responses were suggested to encode an abstract representation of an individual.

Consistent with this hypothesis, recognized famous faces led to greater broadband gamma activity in MTL [55]. However, we found no familiarity-sensitive contacts in amygdala and hippocampus, possibly due to limitations of electrode placements that resulted in only a small number of electrodes were in amygdala and part of hippocampus in our study. The fact that reference electrodes were placed right next to the amygdala in most patients may have also played a role in the lack of observed familiarity sensitivity in amygdala. In MPC, we found more familiarity-sensitive contacts than face-selective contacts and the enhanced activity evoked by famous faces emerged much later than pVTC. This is consistent with the functional role of MPC in memory recall [55,66,67,71,72] and the speculation that MPC is the most abstract extension of the ventral visual pathway [68,69]. Stronger response to famous faces in MPC might be related to their higher emotional content or due to the repeated presentations of stimulus over a relative long period [67].

Neural communication between face areas in the ventral visual pathway and areas critical for retrieval of person knowledge and emotion has been suggested to support face recognition [63,73,74,77**Error! Reference source not found**.]. A recent nonhuman primate study showed that cortical face patches receive inputs from both pulvinar and amygdala [70]. Besides, structural and functional connectivity between cortical areas involved in face processing and medial temporal lobe memory system were revealed in some fMRI studies in humans [62,63,64,65]. Although amygdala has been postulated to have a top-down effect on fusiform gyrus to facilitate facial expressions representation, the function of these connections remains unclear [74].

Our intracranial recoding results showed enhanced connectivity between cortical face processing areas and AMG/HPC during familiar face processing in multiple bands. Notably, viewing famous faces increased the connectivity between pVTC and AMG/HPC at both early and later stages. The early interaction enhancement emerged during the same time period when pVTC is active for the initial face-specific processing. We suggest that familiar faces might prime the retrieval of names, increase emotional arousal and activate emotional memories through fast interactions between fusiform gyrus and AMG/HPC and bypass the more detailed processing in the ventral temporal pathways to support the quick detection of familiar faces [56]. Future work is needed to elucidate the functional role of the early interactions between pVTC and AMG/HPC.

According to the Bottom-up tunning hypothesis, long-term visual experience with specific faces tunes the existing circuitry for the familiar features and consequently the representation of familiar faces is enhanced [1,26]. Another model proposes that repeated exposure to familiar faces enables efficient retrieval of associated person knowledge, which in turn facilitates familiar face processing in a top-down manner [7]. Our results provide important information on the dynamics for face familiarity effect in temporal lobe and could help to differentiate between these two hypotheses. With the observation of the familiarity induced enhanced signal in posterior temporal cortex, our study lends more support for the top-down hypothesis. Some EEG and MEG studies have suggested that face familiarity enhances N170/M170 and identity and gender representations over the whole brain [26,28,29,30,31,32,33,34,35,36].

However, because of the low spatial resolution of MEG and EEG, it is difficult to know where did these early modulation originate from. One possibility is that a subcortical pathway may become tuned to face familiarity and engaged in familiar face processing early.

While sEEG provides excellent spatial and temporal resolution at the same time, we would like to acknowledge that the electrode coverage in sEEG is limited, especially anterior temporal lobe and occipital lobe as observed in our study. Future studies will need to focus on the differential role of different familiarity-sensitive brain areas and their function of connections between areas during familiar face recognition.

## Methods

### Participants

Data were collected from pre-surgical epilepsy patients with intracranial depth electrodes used to localize seizure focus. All the electrode implantation sites were determined only by the clinical needs. Patients had at least one electrode implanted in temporal lobe were included in the experiment. After giving written informed consent, 20 patients (14 males, 6 females, 14-53 years) participated in the experiments in Guangdong San Jiu Brain Hospital. All experimental procedures were approved by the Ethics Committee of School of Psychology, South China Normal University (# 2020-040-04).

### Electrode Implantation and Recording

The electrodes were implanted using Rosa Surgical Robot (ROSA; Medtech, France). Anatomical 3T MRI scans were acquired before surgery and CT scans were acquired post-implantation. sEEG data were recorded at 2000 Hz sampling rate for 12 patients and 1000 Hz for 8 patients due to the limitation of storage capacity of recording system. The online reference during acquisition is either a scalp electrode (in 4 patients) or the average of two recording contacts in the white matter near amygdala (in 16 patients).

In each subject, contacts in each region of interest (ROI: pVTC, aVTC, pSTC/MTC, aSTC/MTC, MPC, amygdala and hippocampus) were identified according to the subject’s own anatomical landmarks. For group visualization (For example, **Figure 1B**), individual MRI anatomical images were aligned to CT images to localize contact positions, and then they were normalized to the standard template in MNI-space using AFNI. Contacts from all 20 subjects were displayed on fsaverage surface provided by FreeSurfer (http://surfer.nmr.mgh.harvard.edu). The contacts were presented either larger or smaller than their real sizes to make it more convenient to display the results.

### Experiment design

The stimuli were 7°×7° photos of grayscale famous faces (well-known celebrities), unfamiliar faces and objects. They were presented to patients on a laptop display at a viewing distance of 57 cm, controlled using Psychtoolbox [57] and MATLAB. Before the start of experiment, patients were asked to choose recognizable faces from 60 famous face images and those selected images were used in the following experiment. The number of unfamiliar face and object images was matched to famous face images for each patient. Mean luminance and contrast of each stimulus was matched using SHINE toolbox [58].

In session 1, unfamiliar faces serving as oddball targets were occasionally embedded in a sequence of frequent famous faces and subjects were asked to press a button when an unfamiliar face appeared. Each image was presented for 500 ms, followed by 3 s fixation. There were 8-10 oddball trials and 150 famous face trials. Patients can take a short break every 10 trials. Session 2 is similar to session 1 except that famous faces served as occasional oddball targets embedded in unfamiliar faces. In session 3, unfamiliar faces (150 trials) and objects (150 trials) were presented, with faces from another race served as the oddball targets (16-20 trials). The order of sessions was randomized across patients.

### Data analysis

sEEG data analysis was performed using MNE-python [59] and MATLAB.

#### Pre-processing

First, raw data were down-sampled to 1000 Hz and filtered with a 50-Hz notch filter. Bad channels (broken or with more than 50% high amplitude noise) and epochs with epileptiform activity were removed by trial-by-trial visual inspection with the help of a neurologist specialized in epilepsy. To remove signal drifts, the data were high-pass filtered at 2 Hz using a 4th order Butterworth filter. Then the filtered data were rereferenced by subtracting the mean of the remaining contacts.

#### HFB analysis

For each contact in each patient, the following analyses were performed: (1) Pre-processed data were band-pass filtered (110–140 Hz, Butterworth, order 4); (2) Hilbert transform was applied and the analytic amplitude was extracted; (3) Down-sampled to 400 Hz and the results were square-root transformed; (4) normalized the results by z-scoring with respect to all baseline periods (from 300 ms to 0 ms before stimulus onset)[60].

Contacts were defined as “active” if they showed a significantly higher average HFB responses between 100 and 700 ms after stimulus presentation compared to the event baseline for at least one category. Significance was assessed by using Wilcoxon rank-sum tests (p < 0.05, FDR correction by the number of electrodes within each ROI). Out of the task-active sites, “face-selective” contacts were defined as contacts where significantly higher responses were seen to faces compared to objects (non-parametric cluster-based permutation tests, FDR corrected) in the time window [100 700] ms after onset. “Familiarity-sensitive” contacts (famous faces > unfamiliar faces) were identified in similar way. Notice that session 1 and 2 were used to do familiarity-sensitive contacts identification and session 3 was used for face-selective contacts identification.

#### Connectivity analysis

Phase locking value (PLV) [51] was calculated for each task-active contact pair between pVTC-AMG/HPC, aVTC-AMG/HPC, pSTC/MTC-AMG/HPC, aSTC/MTC-AMG/HPC, and MPC-AMG/HPC, for famous faces and unfamiliar faces respectively [61]. Only patients with electrodes implanted in both temporal lobe and MTL were included (n=17). The pre-processed data was filtered in the desired frequency band of interest (3-5 Hz, 8-12 Hz, 12-30 Hz, 30-60 Hz) using a FIR filter (order = 4 cycles of the desired signal). Then Hilbert transform was applied and PLV was calculated for each contact pair and normalized to baseline (−300 to 0 ms). For each ROI and each band, we statistically compared the PLVs between famous and unfamiliar faces using a cluster-based permutation test (N=10000, cluster p <0.05, Bonferroni corrected).

## Funding

The Key-Area Research and Development Program of Guangdong Province 2019B030335001 (QG)

The National Science Foundation of China 20 & ZD296 ()

The Key Project of Science and Technology Program of Guangzhou 201804020085 ()

Key Research Program of Frontier Sciences, CAS KJZD-SW-L08 ()

Strategy Priority Research Program of CAS XDB32020206 ()

Ministry of Science and Technology Major Project 2021ZD0204201 ()

